# The bacterial genetic determinants of *Escherichia coli* capacity to cause bloodstream infections in humans

**DOI:** 10.1101/2022.12.31.522367

**Authors:** Judit Burgaya, Julie Marin, Guilhem Royer, Bénédicte Condamine, Benoit Gachet, Olivier Clermont, Françoise Jaureguy, Charles Burdet, Agnès Lefort, Victoire de Lastours, Erick Denamur, Marco Galardini, François Blanquart, the Colibafi/Septicoli and Coliville groups

**Author notes:** Corresponding author: Erick Denamur. co-first authors (Judit Burgaya, Julie Marin). co-last authors (Erick Denamur, Marco Galardini, François Blanquart). The names of the collaborators are listed in the Acknowledgement section.

## Abstract

*Escherichia coli* is both a highly prevalent commensal and a major opportunistic pathogen causing bloodstream infections (BSI). A systematic analysis characterizing the genomic determinants of extra-intestinal pathogenic vs. commensal isolates in human populations, which could inform mechanisms of pathogenesis, diagnostics, prevention and treatment is still lacking. We used a collection of 1282 BSI and commensal *E. coli* isolates collected in France over a 17-year period (2000-2017) and we compared their pangenomes, genetic backgrounds (phylogroups, STs, O groups), presence of virulence-associated genes (VAGs) and antimicrobial resistance genes, finding significant differences in all comparisons between commensal and BSI isolates. A machine learning linear model trained on all the genetic variants derived from the pangenome and controlling for population structure reveals similar differences in VAGs, discovers new variants associated with pathogenicity (capacity to cause BSI), and accurately classifies BSI vs. commensal strains. Pathogenicity is a highly heritable trait, with up to 69% of the variance explained by bacterial genetic variants. Lastly, complementing our commensal collection with an older collection from 1980, we predict that pathogenicity increased steadily from 23% in 1980 to 46% in 2010. Together our findings imply that *E. coli* exhibit substantial genetic variation contributing to the transition between commensalism and pathogenicity and that this species evolved towards higher pathogenicity.

## Introduction

*Escherichia coli* bloodstream infections (BSI) are severe diseases with an incidence of around 5 × 10^-4^ to 1 × 10^-3^ per person-year in Europe and the United States (1–5) and a mortality ranging from 10 to 30% (5), and may account for a few percents of all deaths in these countries (4). The increase in incidence of BSI (1, 2), the global emergence of multidrug resistance clones such as ST131 (6–9), and the ageing population all make BSI an important and growing public health problem. A better understanding of the bacterial genetic factors determining pathogenicity (the capacity to cause infection) and virulence (the severity of infection) (10) would improve our understanding of pathophysiology and potentially improve stewardship and control policies.

The primary niche of *E. coli* is the gut of vertebrates, especially humans, where it behaves as a commensal (11). BSI are opportunistic infections. Two main routes of infection are described, digestive and urinary, corresponding to two distinct pathophysiologic entities. BSI with a digestive portal of entry are more severe. Host condition and comorbidities affect virulence (12–14). A few bacterial genetic factors affecting virulence have been reported. In a genome-wide association study (GWAS) conducted on 912 patients, no bacterial genetic factor was associated with outcome (death, septic shock, admission to ICU), possibly because of insufficient power. Alternatively, in a murine model of BSI, a GWAS conducted on 370 *Escherichia* strains have shown that the *Yersinia pestis* High Pathogenicity Island (HPI), and two additional groups of genes involved in iron uptake, were associated with a higher probability of mouse death (15).

There is a rich tradition of comparing *E. coli* strains sampled from commensal carriage vs. in infections to reveal the determinants of pathogenicity (16, 17). These studies often do not sequence full genomes, which prevents the control for bacterial population structure and the discovery of new determinants of pathogenicity beyond already established lists of virulence genes. Moreover, many studies compare *E. coli* from stools vs. from infections in the same individuals (16). This design is interesting because it blocks hosts factors. But it may also have limited power to detect variants associated with infections because it conditions on individuals *with* an infection, limiting the possibility of comparison to the diversity of strains present in stools. So far, no studies investigated the bacterial genetic determinants of the capacity to cause an infection by comparing large numbers of whole genome sequences of bacteria sampled from the gut (commensals) vs. sampled from infections. This may be explained in part by the small number of large commensal strain collections (18). Another difficulty is that host factors, such as age or co-morbidities, are important determinants of infection (19), and must be adjusted for as much as possible when comparing strains sampled in the two contexts. Lastly, the increased availability of large whole genome sequence collections from BSI also established that a small number of sequence types, mainly ST131, 73, 95, 69, 10, are involved in the majority of BSI (20). These STs are rich in virulence associated genes (VAGs) encoding adhesins, iron acquisition systems, protectins and toxins (16, 17), but pinpointing potentially causal individual genetic determinants can only be done in a rigorous GWAS. So far, no systematic examination of the bacterial genetic determinants of *E. coli* pathogenicity has been done by comparison with a large commensal collection.

In the present work, we took advantage of two recently published collections of BSI (1) and commensal (18) strains gathered between 2000 and 2017 in France and with their genome sequenced. We compared BSI and commensal strain genomes at three levels: phylogenomic composition, virulence and resistance gene content, and lastly unitig content in a GWAS. Our goal was to compare the diversity of commensal and BSI strains and to identify specific genomic features affecting the propensity to cause BSI, using both a targeted and a hypothesis-free approach.

## Results

### A dataset of 912 BSI and 370 commensal isolates

We compared the genomes of 912 strains from BSI in adults, originating from two prospective multicentric studies (Colibafi in 2005 and Septicoli in 2016-7 (19, 21)) performed in the Paris area, to the genomes of 370 commensal strains gathered from stools of healthy adult subjects in 2000, 2001, 2002, 2010 and 2017 in Brittany and the Paris area. In-hospital death (or at Day 28) was 12.9 and 9.5% in the Colibafi and Septicoli studies, respectively. Most of the BSI were community acquired (79.6 and 54.3% in the two collections, respectively). To avoid biases, all strains were isolated with similar protocols adapted to the sample origin (BSI and commensal) and sequenced in our laboratory using a similar approach (Illumina technology). To reduce the influence of the origin of the different studies we introduced the date of the study as a covariate, encoding it as a binary variable with the studies collected before and in or after 2010. To account for host factors, we additionally included sex and age as binary variables. For age, the variable was recording if the donor/patient was above 60 years old or not. Finally, we also focused on the reported portal of entry of the BSI strains, which has previously been associated with some genetic variants (**Figure 1A**) (22). The two collections had similar distributions of these variables, with the important exception of the proportion of isolates corresponding to older donors/patients, which is higher (69.43%) in the BSI collection (**Figure 1B**).

**Figure 1.**
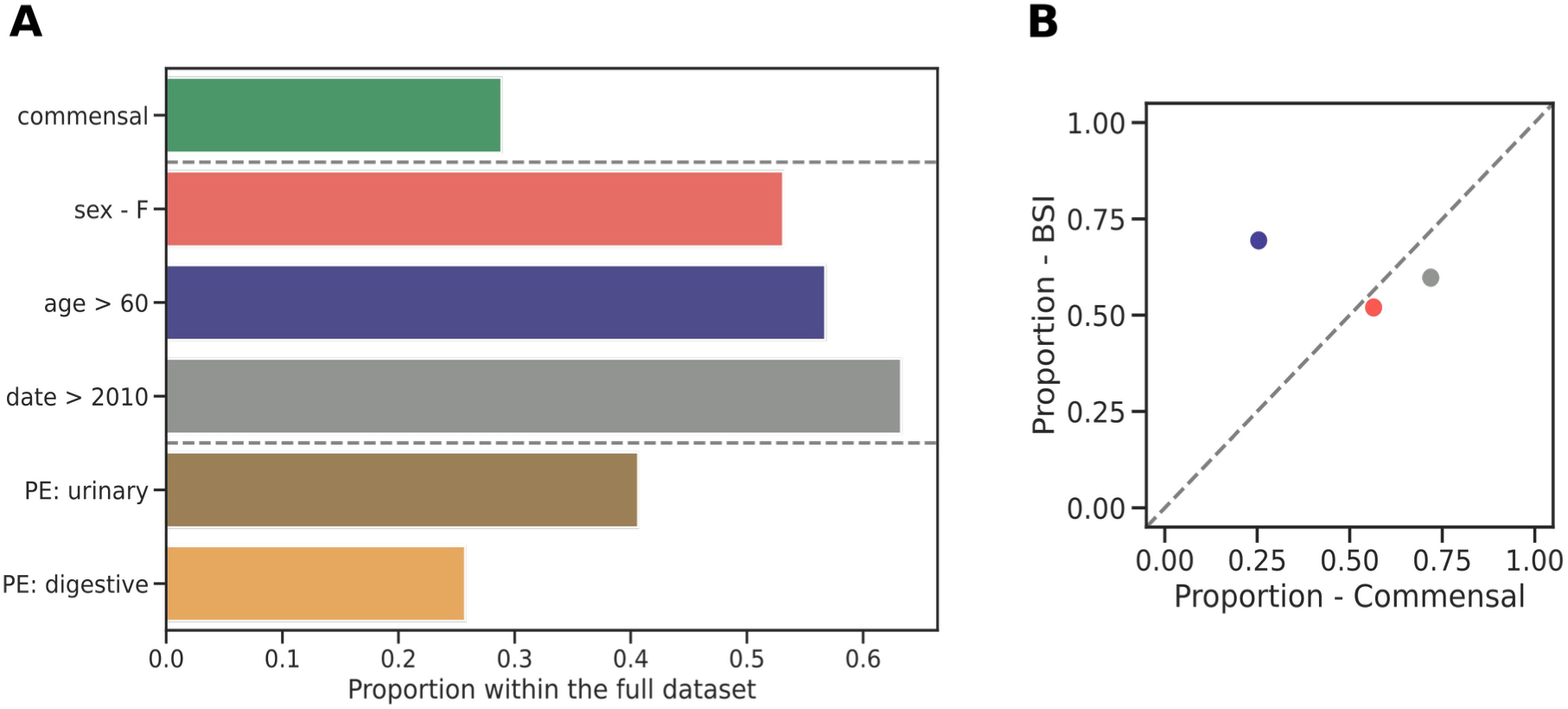
Variables of the combined dataset. (A) Proportion of commensal isolates, distribution of covariates (sex, age, collection date), and BSI isolates with the urinary tract and digestive tract as portal of entry within the full dataset. (B) Scatter plot of the distributions of all covariates in the two collections, colors matching that of panel A. PE: portal of entry.

### Commensal strains are genetically more diverse than BSI strains and have a distinct phylogenetic composition

We first compared the global phylogenomic characteristics of the two collections. The pangenomes of the BSI (N = 912) and commensal (N = 370) collections were composed of 24,321 and 22,373 genes, respectively. For a comparable number of strains, commensal strains had a higher diversity in gene content than BSI strains (**Figure 2A**). Conversely, the core genomes of both collections were similar (3133 and 2985 genes, respectively), and close to the core genome of *E. coli* species as a whole. In terms of SNP diversity of the core genome, the commensal collection was more diverse (pairwise nucleotide diversity π = 2.10e-2) than the BSI collection (π = 2.05e-2, p-value < 2.2e-16).

**Figure 2.**
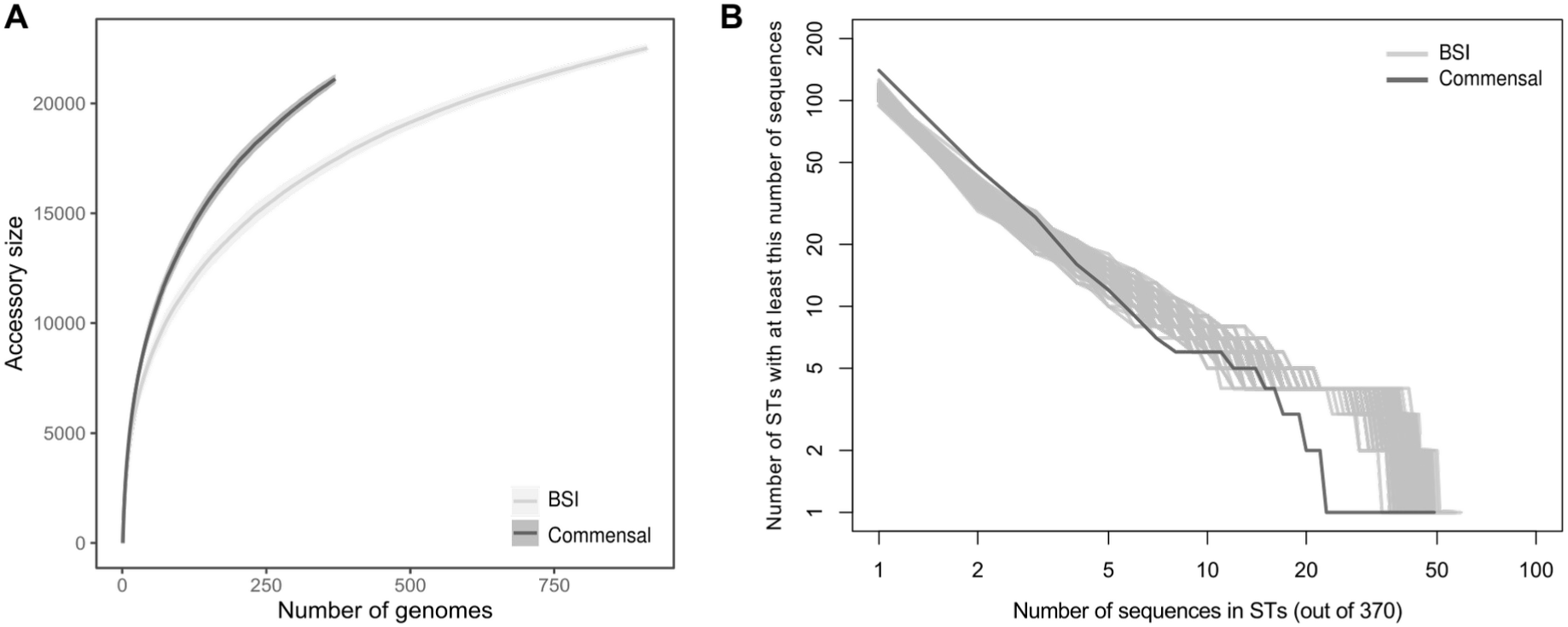
(A) Pangenome sizes as a function of the number of genomes analyzed for the BSI (912 strains) and commensal (370 strains) collections, showing the greater pan genome size of the commensal collection. (B) Cumulative distribution of strain sequences within ST in commensal and BSI collections. To be able to compare the BSI collection with the smaller commensal collection (N = 370), we extracted 200 random sub-samples of 370 sequences from the BSI collection (grey curves).

Commensal strains belong almost equally to A and B2 phylogroups (25.41% and 32.43%) whereas BSI strains belonged mainly to phylogroup B2 (51.21%) followed by D phylogroup (15.79%) (**Table S1**). The commensal collection was more diverse in its ST composition, with a higher number of rare STs and a lower number of frequent STs compared to the BSI collection (**Figure 2B**). This greater phylogenetic diversity could explain both the larger diversity in gene content (23) and larger nucleotidic sequence diversity of the pangenomes of commensals.

As previously noted, the diversity of STs in commensal strains was very distinct to that in BSI strains (**Table S2**). Notably, ST10 and ST59 are abundant in commensal strains (13.2% and 3.8%) but under-represented in BSI strains (3.7% and 0.6%); on the contrary, ST131, ST73, ST69, ST95 are less common in commensal strains than they are in BSI strains. This comparison can be translated in an odds ratio for the risk of infection associated with gut colonization by each ST, which can be seen as a quantitative measure of pathogenicity. The sequence type ST131 is the most pathogenic and ST59 the least pathogenic (**Figure 3** and **Table S2**). When the portal of entry was considered for the ST distribution, a similar pattern was observed for both portals of entry as for the whole collection, although the significance level of the risk of infection might change (**Figure 3** and **Table S2**).

**Figure 3.**
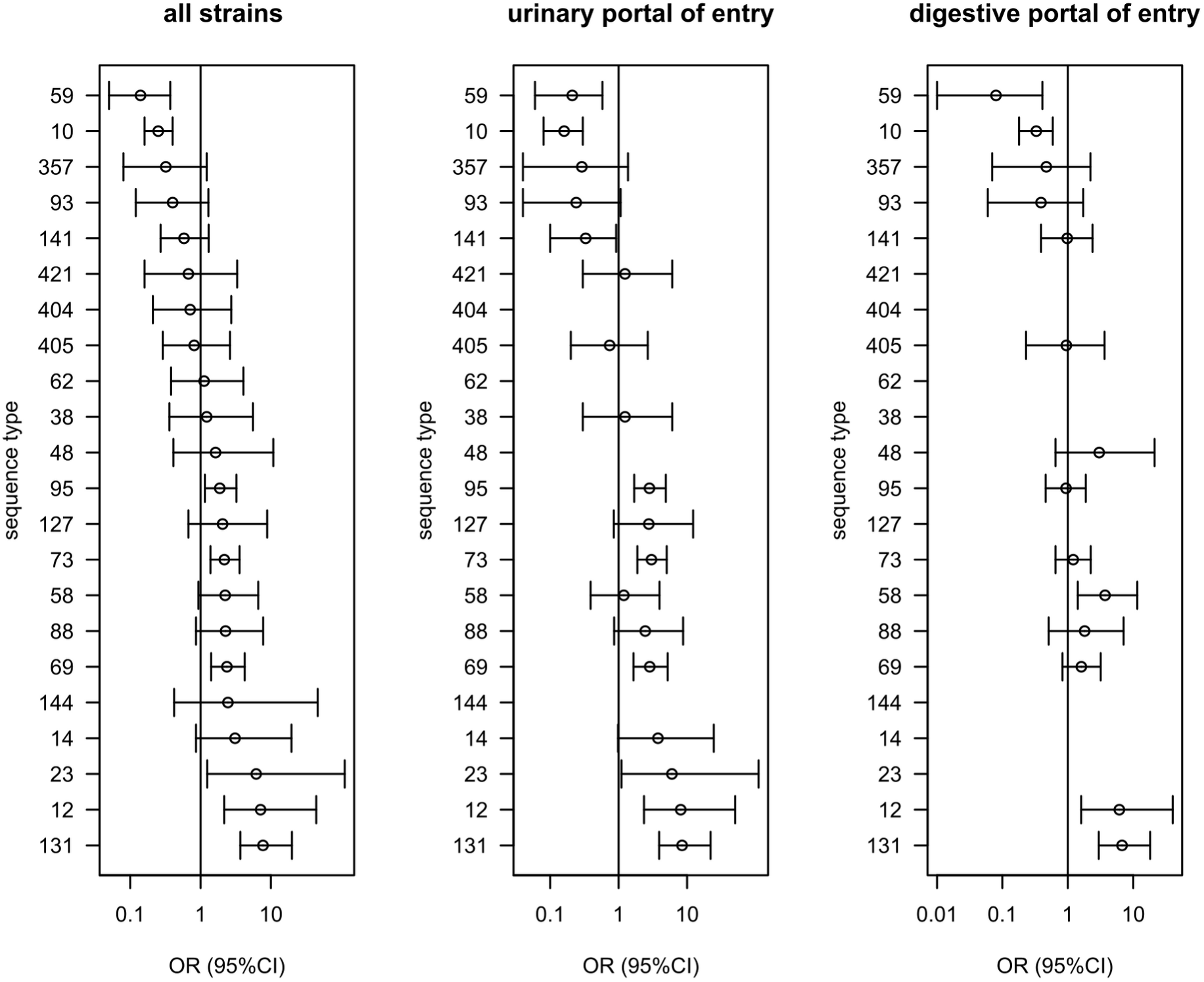
Comparison of the distribution of the sequence types (STs) of the *E. coli* commensal and BSI collections isolates (see table S2). We show the odds ratio (OR with 95% CI) for the risk of infection associated with colonization by each ST (logistic model of infection status as a function of the ST). We selected the STs present in at least 5 strains in at least one of the two collections. STs are ordered by decreasing associated odd ratio for all strains.

The distribution of the O-group diversity also differed between the commensal and the BSI collections (**Table S3**). The four O-groups targeted by the recently developed bioconjugate vaccine ExPEC4V (24, 25), O1, O2, O6 and O25 are the most abundant O-groups in the BSI collection. However, unlike the O-groups O6 and O25, the O-groups O1 and O2 are not particularly associated with BSI strains (**Table S3**). In other words, these two O-groups are frequent in BSI because they are the two most frequent O-groups in commensalism, but are not particularly pathogenic.

### BSI strains are enriched in VAGs and antibiotic resistance genes (ARGs) as compared to commensal strains

Using a targeted approach, we next focused on the frequency of VAGs and ARGs in both collections. A global comparison in the number of VAGs classified in functional categories showed a significantly higher presence of VAGs coding for adhesins, iron acquisition systems, protectins and toxins categories in BSI strains (**Figure 4A and B, Figure S1, Table S5**). We found similar results when comparing against BSI strains with urinary portal of entry to commensals (**Figure 4C**). However, only the iron acquisition systems category remained significant when comparing against BSI strains with digestive portal of entry (**Figure 4D**). More precisely for the full dataset, the highest significance was observed for the *pap* genes with the *papGII* allele, followed by the *sit*, *iuc* and *irp2/fyuA* (HPI) genes, all with p-values << 10^-10^ (**Table S5**). These analyses do not imply a causal role of these genes and alleles in BSI, as they are not adjusted for the distinct phylogenomic composition of commensal and BSI strains. However, it is possible to crudely adjust for this population structure by focusing on the B2 phylogroup strains which are known to exhibit the highest prevalence of VAGs within the *E. coli* species (17).

**Figure 4.**
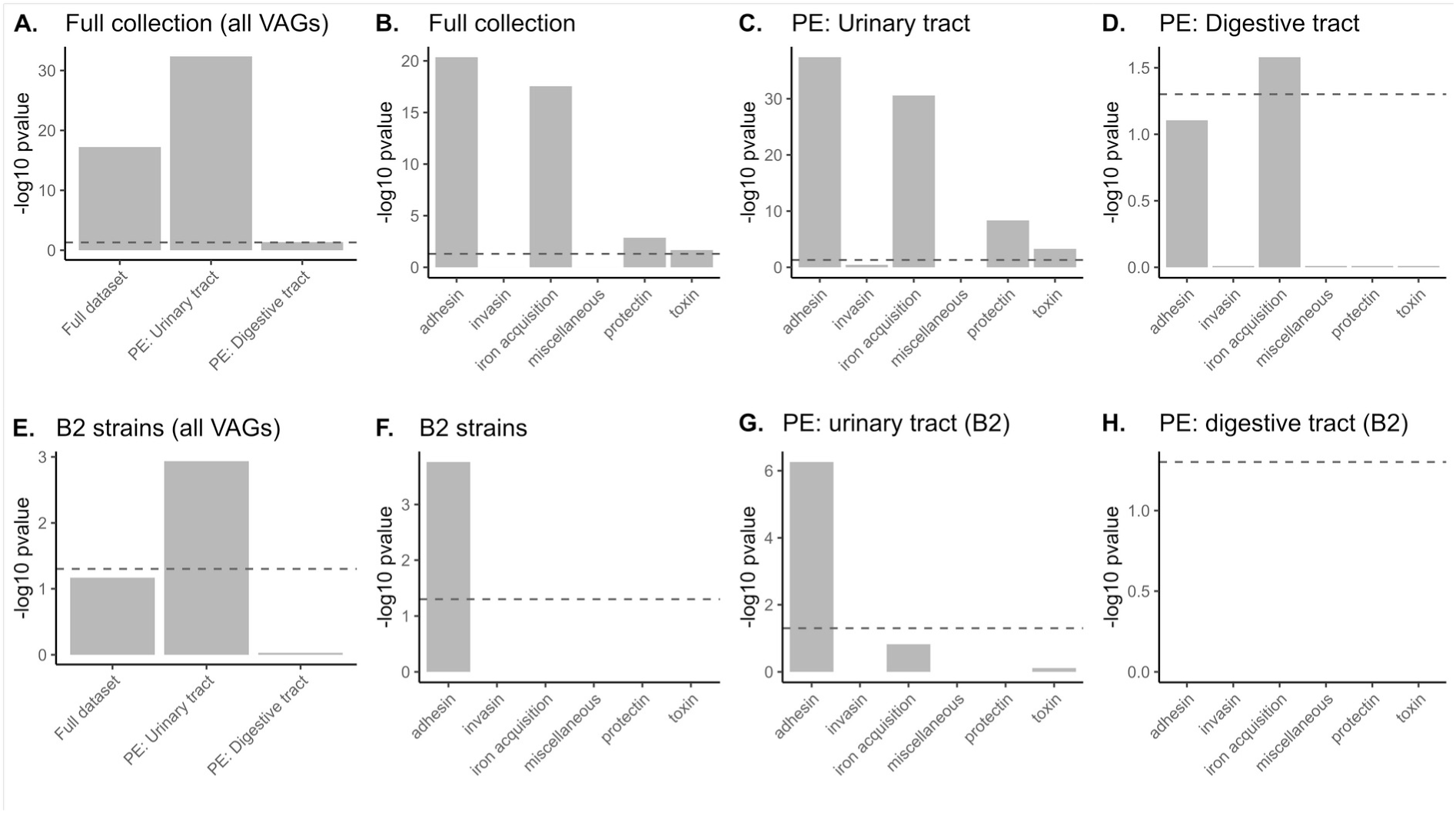
Difference in the number of VAGs per strain among the six main functional classes of virulence between the 912 BSI and 370 commensal strains (Benjamini-Hochberg corrected p value < 0.05). We tested whether the number of VAGs was larger in BSI than in commensal strains considering (A-B) all the strains (912 BSI strains), (C) BSI strains with urinary portal of entry (PE) to commensals (498 BSI strains), (D) BSI strains with digestive portal of entry to commensals (310 BSI strains), (E-F) the B2 strains (467 BSI strains), (G) B2 BSI strains with urinary portal of entry to commensals (304 BSI strains) and (H) B2 BSI strains with digestive portal of entry to commensals (124 BSI strains). The dashed line represents the significance threshold at the 0.05 level.

When only B2 phylogroup strains are compared, only VAGs coding for adhesins category remained significantly over-represented in BSI (**Figure 4F**). When comparing only B2 strains with urinary portal of entry to B2 commensals, again only adhesins were over-represented, and no differences were observed when comparing only against B2 strains with digestive portal of entry (**Figure 4G-H**). Regarding individual genes, interestingly, for two VAGs with experimentally validated role in urinary tract infection, *pap* genes (26) and *fim* genes (27), we found a higher level of significance in B2 strains with urinary portal of entry than in all B2 strains (*pap*) or in all strains (*fimD-H*) (**Table S5**).

BSI strains were predicted to be more resistant to all classes of antibiotics than commensal strains (**Figure 5**). The only exception was for carbapenems (for which resistance was predicted to be very rare). This also holds true if specific portals of entry and/or phylogroup B2 are taken into account (**Figure S2**). To verify that this over-representation of resistance in BSI was not explained by the fact that BSI isolates are slightly more recent on average than commensal isolates, we restricted our analysis to BSI Colibafi strains (sampled in 2005) and found the same results when considering all phylogroups and portals of entry.

**Figure 5.**
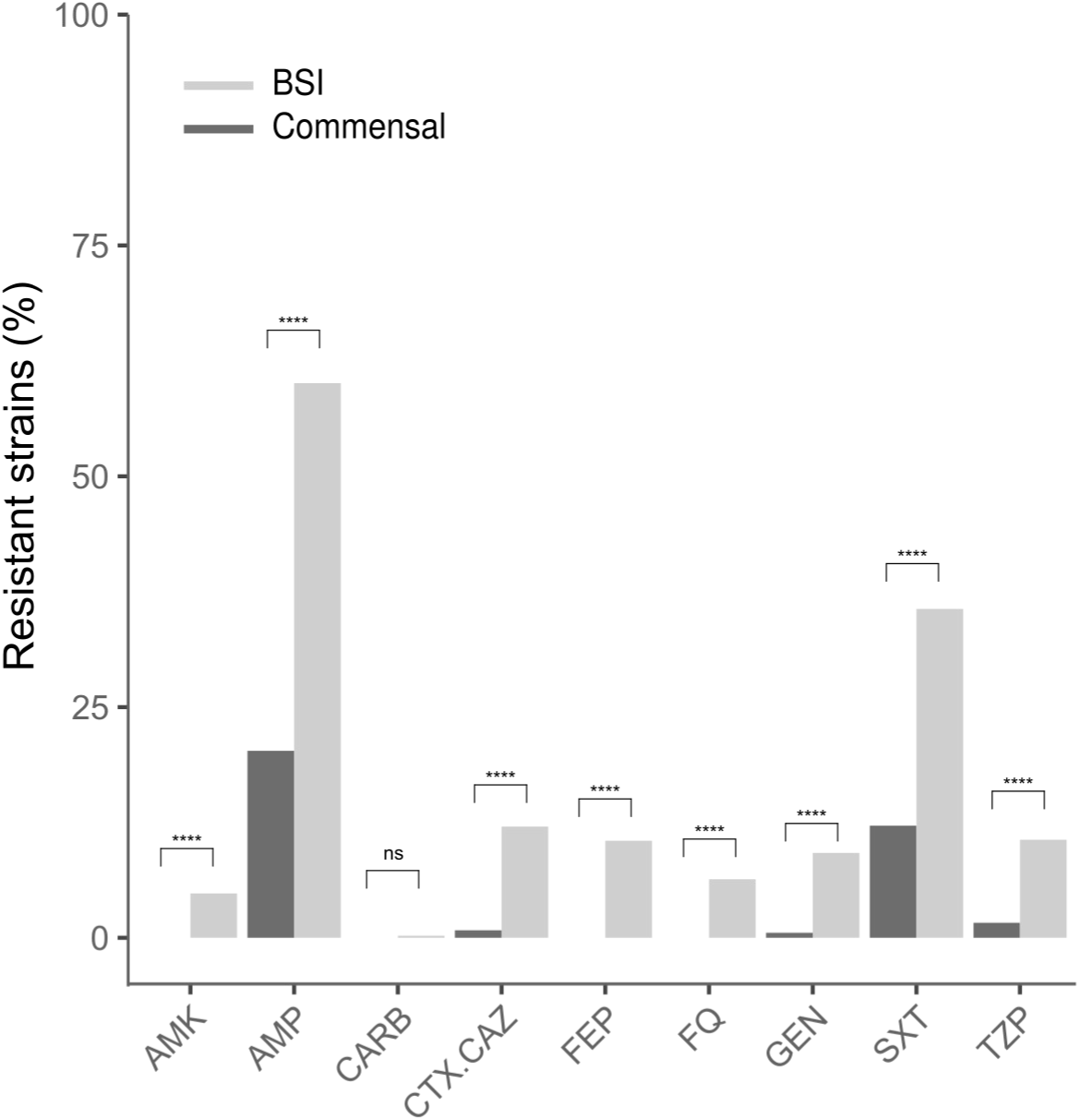
Predicted antibioresistance phenotypes of the 1282 strains (Benjamini-Hochberg corrected p value < 0.05). The results are presented as percentages of resistant strains for nine antibiotics of clinical importance. AMK, amikacin; AMP, ampicillin; CARB, carbapenem; CTX/ CAZ, cefotaxime/ceftazidime; FEP, cefepime; FQ, fluoroquinolones; GEN, gentamicin; SXT, cotrimoxazole; TZP, piperacillin/tazobactam.

No difference in VAG numbers (t-test, all Benjamini-Hochberg corrected p value > 0.05), nor in resistance prevalences (Fisher’s exact test, all Benjamini-Hochberg corrected p value > 0.05), was found when comparing nosocomial and community BSI strains, considering both Septicoli (167 nosocomial and 296 community BSI strains) and Colibafi (75 nosocomial and 292 community BSI strains) collections together or individually.

### Bacterial genetic factors explain a large fraction of the variation in the BSI phenotype

We computed the heritability, as the proportion of the variance of a phenotype explained by variable genetic factors (28), to estimate whether we could expect to find bacterial genetic variants associated with commensalism or BSI in our dataset. We first measured the heritability using the ST information alone, to measure the influence of the genetic background on phenotypic variability. We then computed the heritability emerging from the individual genetic variants (**Figure 6**). We found that STs could explain 24%, 28%, and 11% of the phenotypic variance in the full collection, the subset with BSI isolates with urinary tract as portal of entry and digestive tract as portal of entry, respectively. Genetic variants alone could explain a larger fraction of the phenotypic variability: 65%, 69%, and 39% for the three subsets, respectively. This suggests that pathogenicity might not be solely determined by a strains’ genetic background but also through specific genetic determinants.

**Figure 6.**
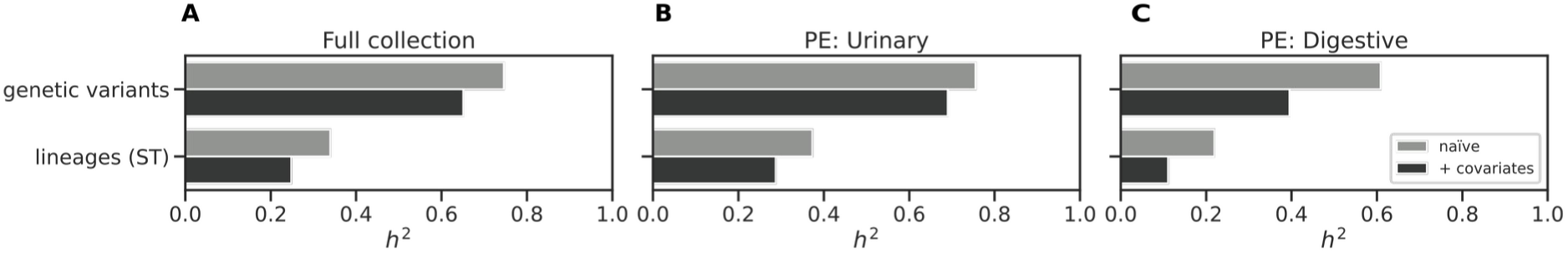
Heritability estimates for the commensal phenotype. A) Heritability estimates for the full dataset. B) Heritability estimates for the subsets with BSI isolates with portal of entry as urinary tract and C) digestive tract.

### A whole-genome machine learning model differentiates commensals from BSI strains

We next applied a machine learning model trained on both the core and accessory genome of the strains to differentiate between commensal and BSI strains and highlight the genetic variants that contribute the most to the discriminatory power of the model (wg-GWAS). We performed the analysis on three different datasets: the full strain collection, and two subsets of BSI isolates: one with urinary tract as portal of entry, and another one with digestive tract as portal of entry. We used all the genetic variants covering the pangenome compactly represented by unitigs and the elastic net linear model implemented in pyseer for the associations (29). We looked for associations between genetic variants and whether a strain was classified as a commensal as the phenotype, and used the following three binary variables as covariates to account for host factors and collection biases: the sex of the donor/patient, their age (older than 60 years old), and the date of each collection (before or after 2010). To quantify model performance, we performed a cross validation by holding out one phylogroup at a time, and computed the precision (proportion of true BSI among the predicted BSI strains), recall (sensitivity) and F1-score (harmonic mean of precision and recall) (**Figure 7** **and S3**). The model performance improved in all cases when the covariates were considered for the associations, potentially confirming that host factors also explain part of bacterial pathogenicity. We also found a better model performance in the two subsets with BSI isolates with a specific portal of entry, compared to the full collection, which could underscore the presence of specific genetic variants associated with either portal of entry.

**Figure 7.**
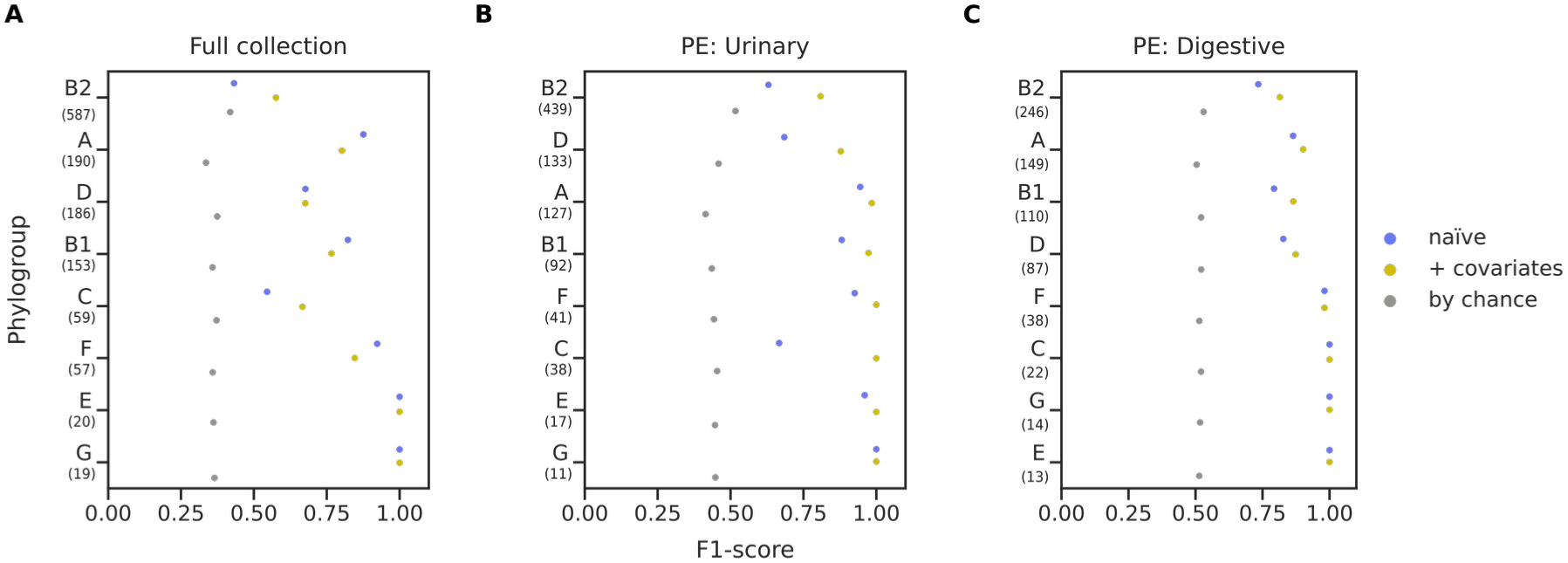
wg-GWAS model performance. F1-score representation. (a) For the full collection (b) the subset of clinical isolates with urinary tract as portal of entry, and (c) the subset of clinical isolates with digestive tract as portal of entry, for the naive analysis (blue dots), with covariates (yellow dots), and the one expected by chance (grey dots). Numbers within parenthesis below each phylogroup indicates the sample size.

We found a number of unitigs to be associated with commensalism (*i.e.* with non-zero weight in the elastic net model). Overall, 107 and 59 unitigs passed the threshold for the model built naïvely and with covariates, respectively, which we then mapped back to 34 and 28 genes. We found that 8 out of the 28 genes obtained through the analysis with covariates were clearly related to virulence. We found the *iucB* gene, encoding an aerobactin siderophore biosynthesis protein (30) and *papG* encoding the adhesin at the tip of the P pilus (31). Both have already been associated with invasive uropathogenic *E. coli* (UPEC) isolates (22, 32). Of note, these genes were identified using the targeted approach after adjusting for population structure by focusing on the B2 phylogroup strains (see above). We also found the following genes: *sopB* which is an inositol phosphate phosphatase associated to virulence in *Salmonella* (33); *mltB*, which is part of a network connecting resistance, membrane homeostasis, biogenesis of pili and fitness in *Acinetobacter baumannii* (34); *fliL*, encoding for the flagellar protein FliL (35). And lastly, two unnamed orthologous groups (group_5900 and group_9261), described as the putative bacterial toxin *ydaT* (36). We found more genes associated to the phenotype when dividing the BSI strains according to their portal of entry. We found a total of 152 and 96 associated unitigs for the urinary and digestive tract subsets, respectively, which we then mapped back to 101 and 45 genes, some of which are known to be involved in pathogenicity and antimicrobial resistance (**Table 1 and S6**). Taken as a whole, we found the associated genes to be enriched in the L COG category (replication, recombination and repair) for the three subsets, and in the K COG category (transcription) for the full dataset only. We also performed a Gene Ontology (GO) term enrichment analysis and found that for the subset with BSI isolates with urinary tract as portal of entry, the relevant (depth > 1) enriched GO terms include different categories related to metabolic processes, ion binding and intracellular anatomical structure (**Table S7**). Similarly, to the targeted analysis described above, we found that the genes resulting from the three associations were enriched for VAGs and ARGs (**Figure 8**); when considering all VAGs and ARGs together we found a significant (pvalue < 0.05) enrichment for the full dataset and the urinary tract subset. We found VAGs related to iron acquisition to be enriched in all three datasets, while adhesins were enriched in the full dataset only. For the ARGs, only the resistance to cotrimoxazole (dfrA for SXT resistance) was enriched in the urinary tract subset.

**Figure 8.**
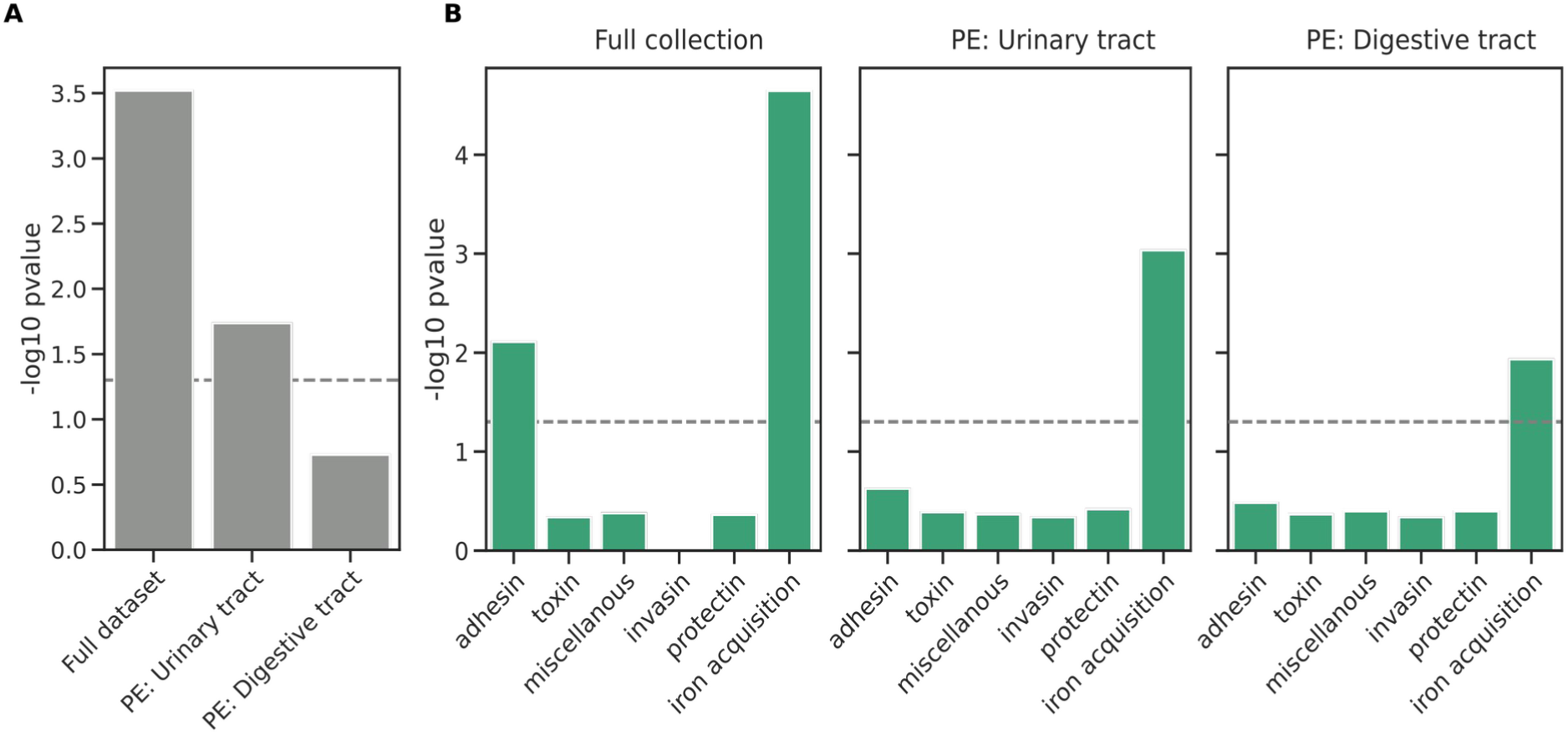
A) Virulence associated genes enrichment analysis for the full set of genes and B) the different functional categories. The significance threshold is represented over the dotted line (Fisher’s exact test, p<0.05). PE: portal of entry.

**Table 1.**
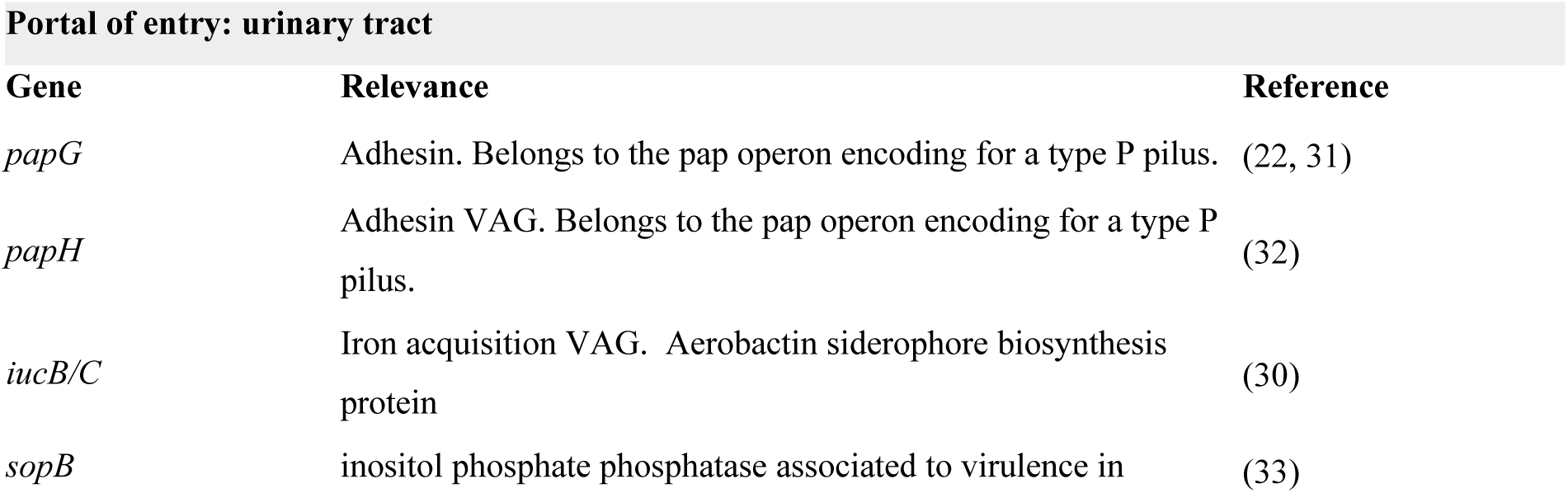
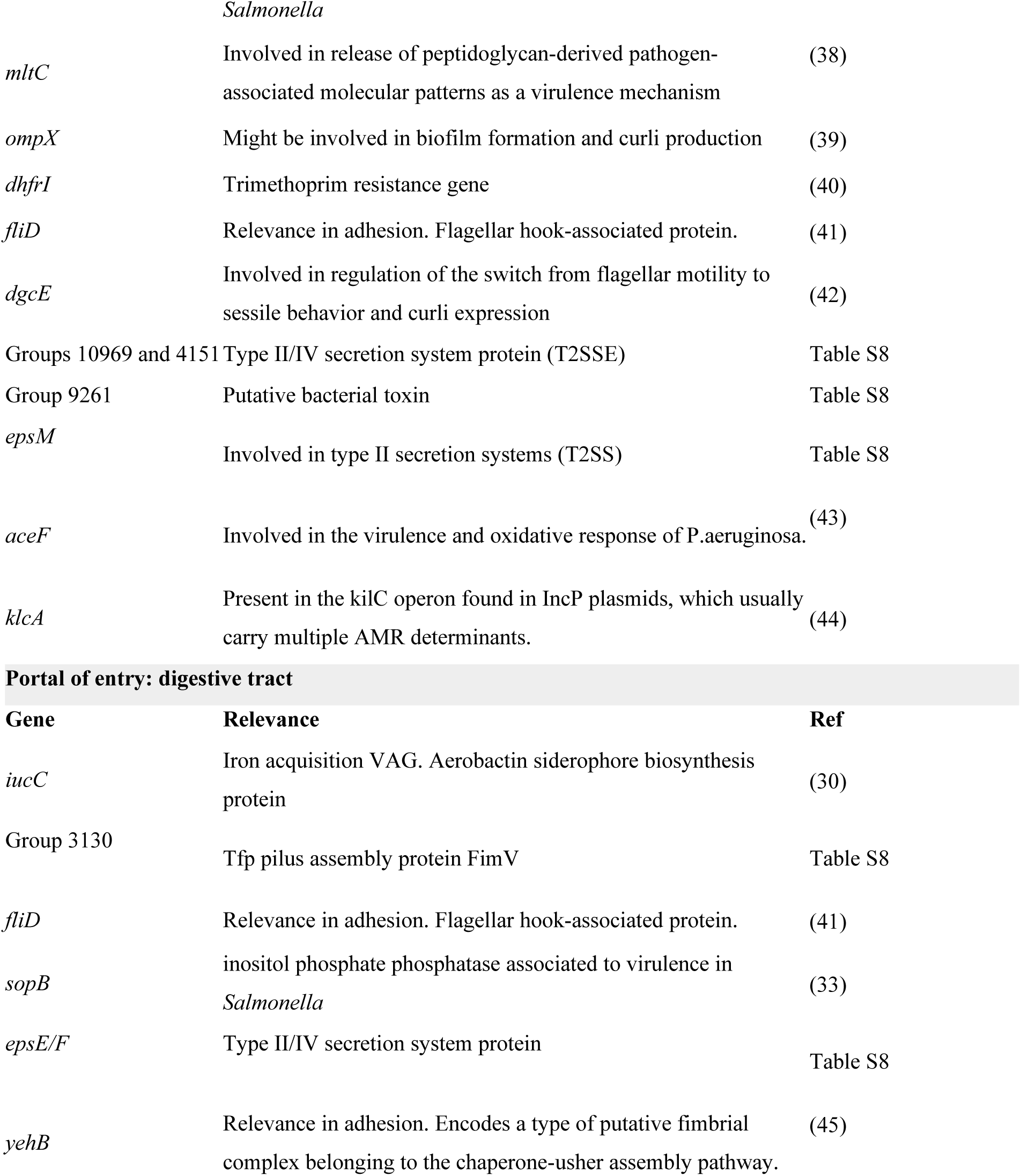
Genes with functions related to pathogenicity and antimicrobial resistance with unitigs associated with the phenotype mapped to them for the two subsets.

The model can be used to predict the potential pathogenicity of other isolates based on the presence of the unitigs for which the model’s weight is different than zero. We predicted the pathogenicity of commensal strains collected at three time periods: 1980 (37), 2000-2002 and 2010. Interestingly, the model predicts a marked increase in pathogenicity of these commensal isolates, with the proportion doubling between the 1980s and the 2010s (23% vs. 46%, **Figure S4**). This suggests that the commensal strains inhabiting the gut of healthy humans may have evolved towards higher pathogenicity in the past decades.

Through an unbiased approach based on the whole pangenome, we have drawn similar results as a more targeted approach, namely that VAGs are to some extent able to distinguish commensals from pathogenic isolates.

## Discussion

It is known since the 1940s (46) that within the *E. coli* species, some strains with a specific genetic background have higher capacity to cause extra-intestinal diseases. Later on, pathogenicity has been associated with specific serotypes, STs, and the phylogroup B2, which are enriched in some VAGs (47, 48). However, disentangling the respective roles of causal genetic variants from the genetic background in a mostly clonal species is a difficult task (49). To do so, we systematically investigated the genomic differences between 912 *E. coli* strains from bloodstream infections and 370 strains sampled from the stools of healthy volunteers.

We revealed differences at three levels. First, at the phylogenetic level, strains from BSI are less diverse, dominated by a small number of highly pathogenic STs, and have consequently smaller pangenomes and lower genetic diversity than commensal strains. Second, strains from infections are enriched in VAGs, and are predicted to be more antibiotic resistant. Third, in a machine learning assisted GWAS, we found 101 and 45 genes associated with BSI with urinary and digestive portal of entry, respectively, independently of the clonal background. Some of these are involved in adhesion and in iron acquisition, as well as other functions. Generally, genes with a significant association are enriched in iron acquisition system, the L COG category (replication, recombination and repair) and GO terms including different categories related to metabolic processes, ion binding and intracellular anatomical structure. The heritability of pathogenicity is estimated at 69% (urinary PE) and 39% (digestive PE), in agreement with the higher role of the host factors in BSI with digestive PE (19, 21). Thus, a large fraction of pathogenicity is explained by bacterial genetic factors. This is roughly double of the heritability when considering STs alone, suggesting that specific genetic variants at a finer phylogenetic scale than ST are determining pathogenicity. For comparison, age, a host factor strongly associated with BSI, explains 17.6% of the variance. Thus, we conclude that bacterial genetics has a significant role in determining pathogenicity, even after basic host factors (age and sex) have been accounted for. An important limitation of our study is that we did not use available information on host co-morbidities in BSI patients for the comparison with commensal strains. In fact, the most frequent co-morbidity in the BSI collection is immunosuppression, which was an exclusion criterion for the commensal collection. Co-morbidities are associated with BSI (5, 18, 22). It is possible that co-morbidities act as a confounder in our study, if they both increase the probability of BSI and influence the *E. coli* strains carried by individuals. If this is the case, the variants we identify may not be directly causal for infections. Rather, they may be bacterial variants that favor the colonization of individuals with co-morbidities. Age is also associated with BSI (5, 14, 50). In this work, we do control for age, albeit in a crude way, with the covariate “above or below 60 years old”. If some of the variation associated with age is not captured by this covariate, some of the variants we identify could favor the colonization of older or younger individuals. For example, there is evidence of age-associated variants in *Streptococcus pneumoniae* (51). To attenuate these concerns on confounding, we remind that several of the significant variants have an experimentally validated role in infection and virulence (**Table 1**).

We found that strains from infections are more likely to be resistant to antimicrobials. What is the mechanism behind this association, also found in similar GWAS conducted on other pathogens (52, 53)? Confounding is a first possibility: hosts with co-morbidities are more likely to develop a BSI and to use antibiotics frequently. Individuals may even be already treated by antibiotics at the time of infection, in which case only resistant strains would be able to cause this infection. If this mechanism operates, we could expect resistance to be more frequent in hospital-associated than in community-associated BSI, if hosts in hospitals are more likely to use antibiotics at the time of infection. However, we did not find any difference between resistance in hospital-associated and community-associated BSI. Second, antimicrobial resistance genes may have a causal role in infection. This seems unlikely given their very specific function. Third, there might be a genetic association (linkage disequilibrium) between resistance genes and genetic determinants of infection (52, 54). In the third case, we expect the association to disappear when controlling for population structure. With this control, we find that indeed, only one out of nine categories of resistance is significantly enriched in BSI compared to commensals. This suggests that antibiotic resistance genes are genetically linked with pathogenicity determinants, and opens the interesting possibility that antibiotic resistance coevolves with pathogenicity determinants associated with the clonal background of *E. coli*.

The present study compares *E. coli* whole genomes in colonization and in infection, as done before for *Klebsellia pneumoniae* (52), *S. pneumoniae* (55), *Staphylococcus aureus* (53, 56), *Neisseria meningitidi*s (57). These GWAS studies presented a range of results, from low heritability (2.6% for *S. aureus* carriage vs. BSI (53)), to intermediate (36.5% for *N. meningitidis* carriage vs. invasive meningococcal disease), and an analogously large heritability of 70% for *S. pneumoniae* invasive disease vs. carriage, along with a handful of significant SNPs (55). We find a large heritability for *E. coli* BSI vs. colonization, which suggests that a vaccine targeted at virulence determinants could reduce (at least temporarily) the burden of infection (24).

The large heritability of *E. coli* capacity to cause infection also implies that this trait can readily evolve. Evolution of *E. coli* pathogenicity would have important public health implications, given that *E. coli* BSI are a major cause of morbidity and death in Western countries. To investigate temporal trends in pathogenicity, we computed the pathogenicity score with the machine learning model (used to predict the commensal vs. BSI status of strains), in a dataset of commensals from 1980 to 2010 in France (18). We found that the proportion of commensal *E. coli* isolates predicted to be pathogenic isolates with our trained model increased over time, from 23% in the collection from 1980 to 46% in the collection from 2010 (Figure S4). Applying this predictive model to the large collection of available *E. coli* genome sequences, which currently numbers to more than 200,000 genomes (58), could unravel the dynamics of pathogenicity across time and space. This effort would however need to be properly controlled for the biases in the isolates sampled and sequenced (most of them coming from infections), and the phylogroup-specific performance of the model.

What selective pressures might act on pathogenicity determinants? The capacity to cause infection may not be selected *per se*, as infections are a relatively rare occurrence in the life cycle of *E. coli* and do not obviously confer a transmission advantage. Pathogenicity determinants have diverse functions and may therefore be selected for a variety of reasons. They may for example improve the ability to colonize the human gut, improve the ability to compete and replace existing strains, or allow longer persistence in the gut (59–62). Elucidating the selective pressures acting on these determinants is an important research question that would improve our understanding of *E. coli* pathogenicity.

This work opens perspectives to improve studies of the determinants of *E. coli* pathogenicity. It remains difficult to pinpoint individual variants because of the clonal structure of *E. coli*, and confounding by host factors is a concern. One idea to alleviate clonal structure is to focus on specific STs. This would limit the dominant effect of STs belonging to phylogroup B2 and carrying many virulence genes. However, the genetic diversity within a single ST might also be limited. This makes it difficult to anticipate the results of such ST-focused studies. Another idea is to extend to whole genomes the line of work comparing strains from infections vs. colonization in the same individuals. This design would block host effects but, as stated in the introduction, implies that power is contingent on the within-host diversity of strains present in colonization. Further help will also likely come from linking pathogen diversity to clinical and epidemiological phenotypes and including the genetic variation of the host into the association such as in a previous study of *S. pneumoniae* (55). Lastly, similar studies should be conducted in low and middle income countries, where a potentially very distinct diversity of *E. coli* circulates (11) and where the public health problem posed by BSI will escalate with the ageing population in the years to come.

In conclusion, we elucidated the bacterial genetic determinants of pathogenicity of the major human pathogen *E. coli*. The capacity to cause BSI, particularly with urinary PE, is strongly determined by sequence types, additional genetic factors, and tens of specific variants. This implies that *E. coli* pathogenicity may evolve, informs future studies of *E. coli* mechanisms of pathogenicity, and opens the possibility to reduce the burden of *E. coli* with a vaccine targeted at these variants.

## Material and methods

### Strain collections

We studied the whole genomes of 1282 *E. coli* strains divided in two datasets, 370 commensals strains and 912 BSI strains. Commensal strains were gathered from stools of 370 healthy adults living in the Paris area or Brittany (both locations in the North of France) between 2000 to 2017. These strains come from five previously published collections: ROAR in 2000 (n=50)(63) (Britanny), LBC in 2001 (n=27)(64) (Britanny), PAR in 2002 (n=27)(64) (Paris area), Coliville in 2010 (n=246)(65) (Paris area) and CEREMI in 2017 (n=20)(66) (Paris area) (**Table S5**). In addition, a collection of 53 commensal strains from 53 healthy subjects in Paris (37) was used to assess the temporal trend of pathogenicity. BSI isolates (Colibafi (n=367) and Septicoli (n=545) collections) were collected at years 2005 and 2016-2017, respectively (67). In all studies, one single *E. coli* colony randomly picked was retained per individual after plating the blood cultures or the stools.

All multicenter clinical trials were approved by the appropriate ethic committees. The Colibafi study was approved by the French Comité de Protection des Personnes of Hôpital Saint-Louis, Paris, France (approval #2004-06, June 2004). The Septicoli study was approved by the French Comité de Protection des Personnes Ile de France n°IV (IRB 00003835, March 2016). Because of their non-interventional nature, only an oral consent from patients was requested under French Law. The study on the commensal strains was approved by the ethics evaluation committee of Institut National de la Santé et de la Recherche Médicale (INSERM) (CCTIRS no. 09.243, CNIL no. 909277, and CQI no. 01-014).

All the sequences were available (Bioproject PRJEB38489 (ROAR), PRJEB44819 (LBC), PRJEB44872 (PAR), PRJEB39252 (Coliville), PRJEB39260 (Colibafi) and PRJEB35745 (Septicoli)) except the 20 samples of the CEREMI collection that were whole-genome sequenced in the present work, following the protocol detailed in (21) (Bioproject PRJEB55584).

### Genomic diversity of the core genome

The 1282 assemblies were annotated with Prokka v1.14.6 (68). We then performed pan-genome analysis from annotated assemblies with Panaroo v1.3.0 with strict clean mode and the removal of invalid genes (69). We generated a core genome alignment spanning the whole set of core genes as determined by Panaroo, and a phylogenetic tree was computed using FastTree v2.1.11 (70).

### Comparison of commensal and BSI E. coli collection

Multilocus sequence typing (MLST) was performed using an in-house script Petanc, that integrates several existing bacterial genomic tools (71). We determined STs (Warwick MLST scheme) (72) and O types (73).

We evaluated the risk of infection associated to colonization by a specific ST and by a specific O-group. We compared the ST and O-group diversity from the collection of 912 BSI isolates with the 370 commensal isolates, for all STs with at least 5 strains in at least one of the two collections and for all O-groups with at least 5 strains in at least one of the two collections.

The odds ratios for the infection risk were computed by fitting a logistic model of infection status (commensal or BSI) as a function of the ST or the O-group (here and thereafter, “significant” refers to significance at the 0.05 level).

Next, we compared the phylogenetic distribution of the commensal collection with the BSI collection. For all strains, we calculated the cumulative frequency distribution of STs in the commensal collection, and we compared it to the same distribution in 200 random sub-samples of 370 sequences from the BSI collection.

We plotted the pangenome variation with the number of genomes analyzed (Panaroo output). We evaluated the pangenome variation between commensal and BSI isolates with Panstripe (74) using the output of FastTree (phylogeny of all strains) and Panaroo (gene presence absence matrix). We randomly subsampled 100 trees of 370 tips from the BSI phylogeny (n=912) and compared the rate of gene gain and loss between those trees and the commensal tree (n=370). To quantify the genetic diversity, we computed the pairwise nucleotide diversity (π)(75) in R (package ape)(76).

We also compared the number of virulence factors and the proportion of resistance strains between commensal and BSI isolates. We performed t-tests to compare the distribution of VAGs for each of the six main functional classes (adhesin, invasin, iron acquisition, miscellaneous, protectin and toxin) and reported effect sizes using Cohen’s d. Next, we performed Fisher’s exact tests to compare the proportions of strains carrying each VAG of a given functional class between commensal and BSI isolates. All p-values were corrected for multiple comparisons with the Benjamini-Hochberg method, with a 5% family-wise error rate.

We predicted phenotypic resistance as described in (67) for nine antibiotics of clinical importance (amikacin, ampicillin, carbapenem, cefotaxime/ceftazidime, cefepime, fluoroquinolones, gentamicin, cotrimoxazole and piperacillin/tazobactam). We compared the distribution of strains predicted to be resistant on each of the nine antibiotics using Fisher’s exact tests. We again corrected the p-values for multiple tests with the Benjamini-Hochberg method.

### Heritability estimates

We estimated narrow-sense heritability for the target variable using 2 different covariance matrices: one built from the phylogroup using a kinship matrix, and another one with the age. Limix v3.04 (77) was used, assuming normal errors for the point estimate.

### Association analysis

We derived unitigs using unitig-counter v1.1.0 (78). We tested locus effects using the wg (whole genome) model of pyseer v1.3.6 (29, 79), which trains a linear model with elastic net regularization using the presence/absence patterns of all unitigs. We used an alpha with value of 1 for the elastic net, which is equivalent to a lasso model. Cross-validation was performed by holding out each phylogroup sequentially. The model performance was assessed by computing three metrics using each phylogroup. The precision, as the measure of how many positive predictions made are correct; the recall, as the measure of how many positive cases the classifier predicted correctly over all the positive cases; and the F1-score, as the harmonic mean of the two metrics. The F1-score expected by chance was computed overall, for each phylogroup and for the different subsets, by randomly assigning the phenotype to the test samples and running 1000 randomizations. The unitigs with a non-zero model coefficient were mapped back to all input genomes, and gene families were annotated by taking a representative protein sequence from all genomes encoding each gene, which was then used as the input for eggnog-mapper v2.1.3 using the panaroo output to collapse gene hits to individual groups of orthologs. GO terms enrichment was determined using goatools v1.2.3 (80). An *in-house* list of *E. coli* virulence genes and antibiotic resistance genes was used to annotate the virulence and antibiotic resistant genes within the collection, and a Fisher’s exact test was used to determine the enriched genes, with a multiple testing correction based on the Benjamini-Hochberg method, with a 5% family-wise error rate. For the COG and virulence genes enrichment analysis a random ST131 genome from the full dataset was picked up as background.

### Prediction analysis

We used unitig-caller v1.3.0 (81) to make variant calls in the test population, and the elastic net regularization, previously trained, model using pyseer v1.3.6 (79) to predict the phenotype in new commensal samples from different time periods, divided in decades.

### Code availability

Apart from the software packages mentioned in the previous sections, the following were used to run the analysis and generate the visualizations presented in this work: pandas v1.3.4 (82), numpy v1.20.3 (82), scipy v1.7.1 (83), matplotlib v3.4.3 (84), seaborn v0.11.2 (85), biopython v1.80 (86) jupyterlab v3.2.1 (87). Most of the analysis were incorporated in a reproducible pipeline using snakemake v7.18.1 (88) and conda v4.10.3 (89), which is available as a code repository on GitHub (https://github.com/jburgaya/2022_ecoli_commensal) under a permissive licence (MIT).

## Acknowledgments

ED was partially supported by the “Fondation pour la Recherche Médicale” (Equipe FRM 2016, grant number DEQ20161136698). GR was supported by a “Poste d’accueil” funded by the “Assistance Publique-Hôpitaux de Paris” (AP-HP) and the “Commissariat à l’énergie atomique et aux énergies alternatives” (CEA) personal grant for his PhD. MG and JB were supported by the Deutsche Forschungsgemeinschaft (DFG, German Research Foundation) under Germany’s Excellence Strategy - EXC 2155 - project number 390874280. JB was further supported by the Deutsche Forschungsgemeinschaft grant number GA 3191/1-1.

The Colibafi/Septicoli group is composed of: Michel Wolff, Loubna Alavoine, Xavier Duval, David Skurnik, Paul-Louis Woerther, Antoine Andremont, Etienne Carbonnelle, Olivier Lortholary, Xavier Nassif, Sophie Abgrall, Françoise Jaureguy, Bertrand Picard, Véronique Houdouin, Yannick Aujard, Stéphane Bonacorsi, Agnès Meybeck, Guilène Barnaud, Catherine Branger, Agnès Lefort, Bruno Fantin, Claire Bellier, Frédéric Bert, Marie-Hélène Nicolas-Chanoine, Bernard Page, Julie Cremniter, Jean-Louis Gaillard, Françoise Leturdu, Jean-Pierre Sollet, Gaëtan Plantefève, Xavière Panhard, France Mentré, Estelle Marcault, Florence Tubach, Virginie Zarrouk, Frederic Bert, Marion Duprilot, Véronique Leflon-Guibout, Naouale Maataoui, Laurence Armand, Liem Luong Nguyen, Rocco Collarino, Anne-Lise Munier, Hervé Jacquier, Emmanuel Lecorché, Laetitia Coutte, Camille Gomart, Ousser Ahmed Fateh, Luce Landraud, Jonathan Messika, Elisabeth Aslangul, Magdalena Gerin, Alexandre Bleibtreu, Mathilde Lescat, Violaine Walewski, Frederic Mechaï, Marion Dollat, Anne-Claire Maherault, Mélanie Mercier-Darty, Bernadette Basse, Bruno Fantin, Xavier Duval, Etienne Carbonnelle, Jean-Winoc Decousser, Raphaël Lepeule.

The COLIVILLE group is composed of: Monique Allouche, Jean-Pierre Aubert, Isabelle Aubin, Ghislaine Audran, Dan Baruch, Philippe Birembaux, Max Budowski, Emilie Chemla, Alain Eddi, Marc Frarier, Eric Galam, Julien Gelly, Serge Joly, Jean-François Millet, Michel Nougairede, Nadja Pillon, Guy Septavaux, Catherine Szwebel, Philippe Vellard, Raymond Wakim, Xavier Watelet and Philippe Zerr.

---

**Figure S1.**
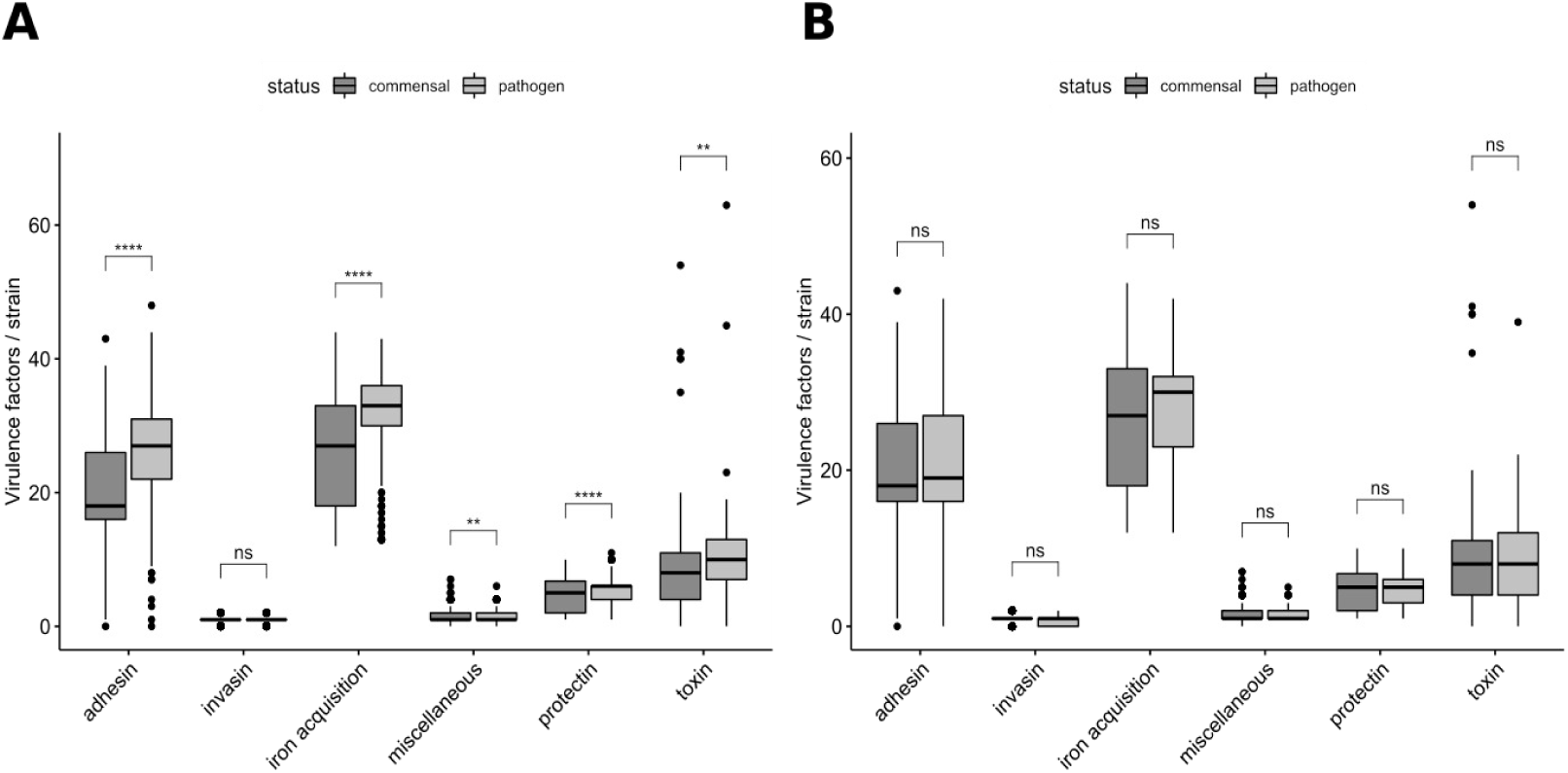
Distribution of the number of virulence factors per strain among the six main functional classes of virulence (Benjamini-Hochberg corrected p value < 0.05) for (A) all the strains with a urinary portal of entry and for (B) all the strains with a digestive portal of entry. Significant differences are indicated by asterisks (p value < 0.05: *; p value < 0.01: **; p value < 0.001: ***; p value < 0.0001: ****; ns: non-significant).

**Figure S2.**
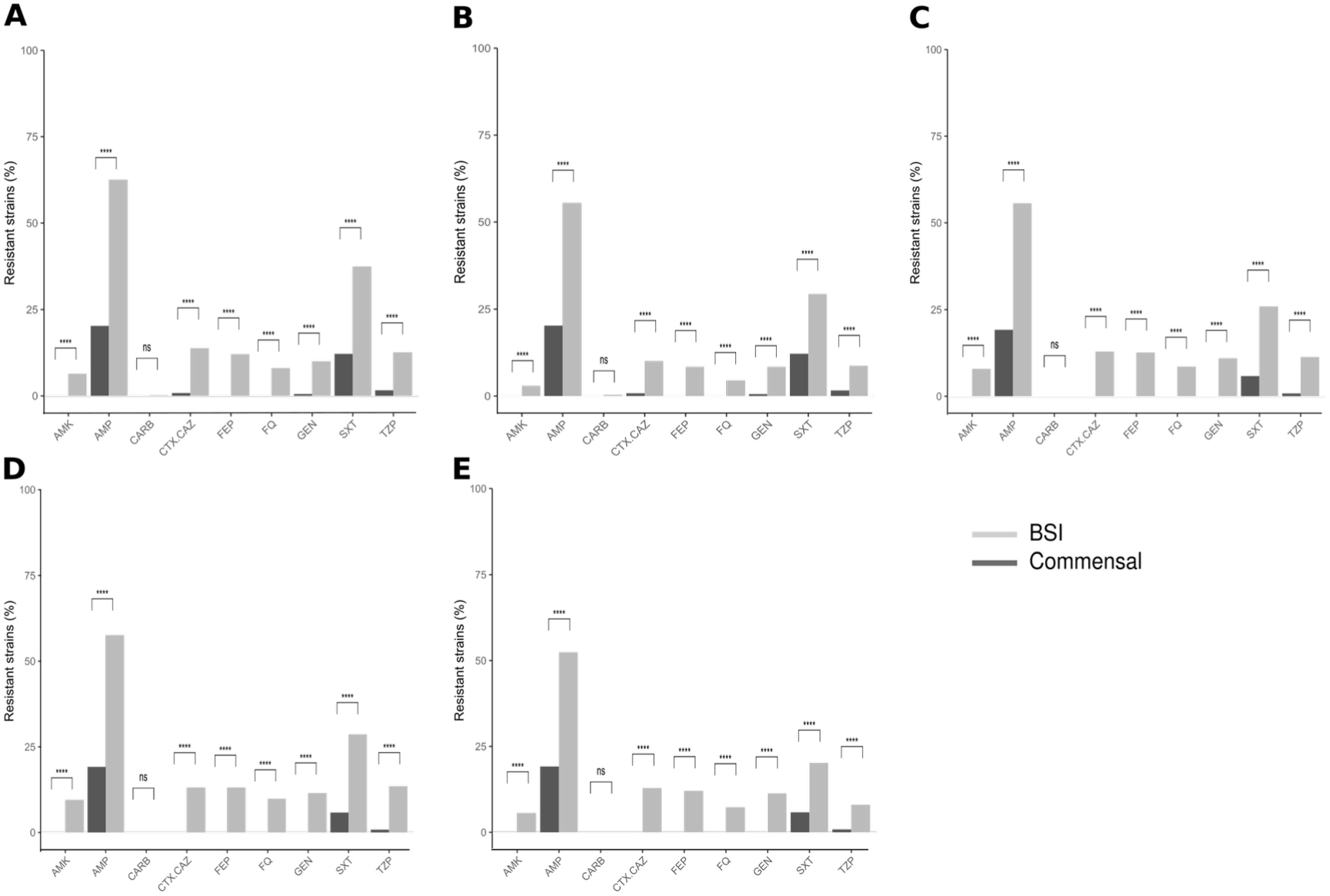
Predicted antibioresistance phenotypes of A) all the strains with a urinary portal of entry, (B) all the strains with a digestive portal of entry, C) B2 strains, D) B2 strains with a urinary portal of entry and (E) B2 strains with a digestive portal of entry (Benjamini-Hochberg corrected p value < 0.05). The results are presented as percentages of resistant strains for nine antibiotics of clinical importance. AMK, amikacin; AMP, ampicillin; CARB, carbapenem; CTX/ CAZ, cefotaxime/ceftazidime; FEP, cefepime; FQ, fluoroquinolones; GEN, gentamicin; SXT, cotrimoxazole; TZP, piperacillin/tazobactam. Significant differences are indicated by asterisks (p value < 0.0001: ****; ns: non-significant).

**Figure S3.**
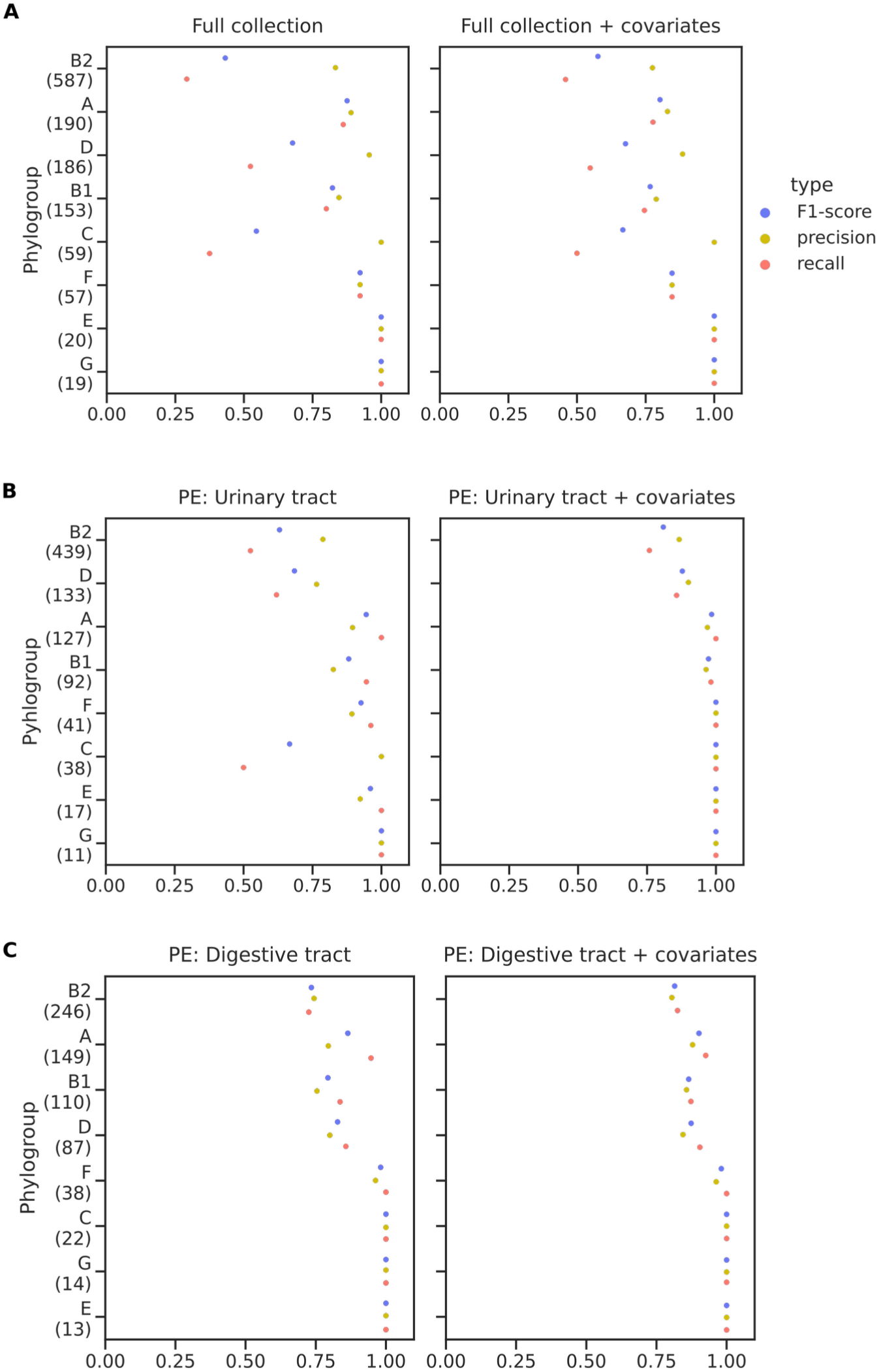
wg-GWAS model performance. F1-score representation (blue dots), precision (yellow dots), and recall (red dots). A) For the full collection B) the subset of clinical isolates with urinary tract as portal of entry, and C) the subset of clinical isolates with digestive tract as portal of entry. The naive and the analysis with covariates are represented. PE: portal of entry.

**Figure S4.**
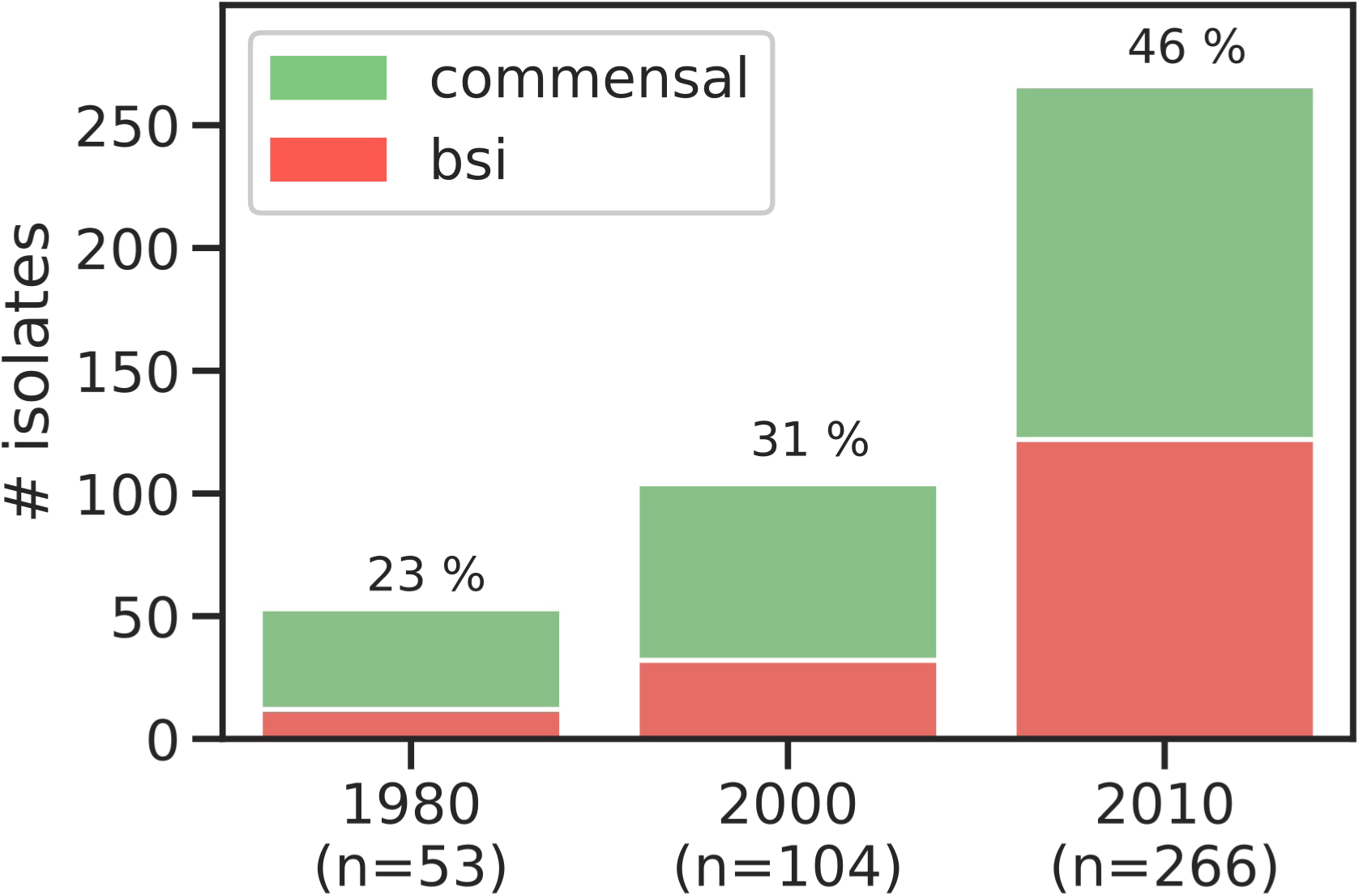
Proportion of BSI predicted isolates over time. 423 isolates from commensal collections were fitted to the trained ML model. The proportion of BSI isolates for the 3 different periods of time is colored in red and the percentage indicated above each bar. The total number of isolates per year is given in brackets.

